# Behavioural, biochemical and functional phenotyping of chronic exposure to chlordecone in mice

**DOI:** 10.1101/2025.06.16.659846

**Authors:** Johann Vulin, Fabien Chauveau, Chloé Dure, Muris Humo, Marco Valdebenito, Claire Aufauvre, Louis Vidal, Eric Morignat, Jérémy Verchere, Aude Decesar, Damien Gaillard, Latifa Lakdhar, Thierry Baron, Gwenaëlle Lavisson-Bompard, Benjamin Vidal

**Author notes:** Corresponding author: Thierry BARON. (co-last authors) (Thierry Baron, Gwenaëlle Lavisson-Bompard, Benjamin Vidal).

## Abstract

**Background:** Chlordecone (CLD) is a persistent organochloride pesticide formerly used against banana weevil. It is detectable in blood samples from a large proportion of the population in the French Caribbean Islands. Several experimental studies have demonstrated acute neurotoxicity of CLD, but the effect of a subchronic exposure to CLD remains to be studied.

**Methods:** Young adult male mice were injected intraperitoneally with 3 mg/kg CLD (n=34) or vehicle (n=22), twice a week, for eight weeks. Behavior, regional brain accumulation, and effects on the dopaminergic system were studied. In addition, functional ultrasound imaging (fUSi) was used to probe the visual, somatosensory and dopaminergic pathways.

**Results:** CLD was detected in all brain regions (5-15 mg/kg) after two-month exposure, without any marked impact on behavior (anxiety, motor coordination, memory). The dopaminergic system was mostly unaffected, despite slight decreases in the number of TH-positive neurons and the expression of VMAT2, quantified in a subset of animals. fUSi highlighted a decreased response to the visual stimulation in CLD-exposed animals, in contrast to the sensorimotor response, which was found unaltered.

**Conclusion:** The two-month-long, systemic, exposure to an intermediate dose of CLD resulted in a mostly unaffected phenotype, with a normal behavior and a largely intact dopaminergic system. Interestingly, functional ultrasound imaging was able to detect an altered visual response, which has also been noted in Parkinson’s disease. This study position functional ultrasound imaging as a promising technique to capture early signs of neurotoxicity, opening up opportunities for “toxico-fUS” in the field of neurotoxicology.

**Highlights:** High CLD neurotropism confirmed in mice by LC-MS/MS.

Sub-chronic chlordecone exposure suggests possible early signs of parkinsonism.

Functional UltraSound reveals impairment of brain areas linked to vision and hearing.

## 1. Introduction

In the French Caribbean Islands of Guadeloupe and Martinique, from 1972 until 1993, the organochloride pesticide chlordecone (CLD) was extensively used against banana weevil, an important pest of banana plantations. Mainly due to the high affinity and retention capacity of organic soils and low solubility in water, CLD is still found in waterways and soil, and consequently in food products of plant and animal origin, including seafood products. This explains why CLD is detectable in blood samples from a large proportion of the population of the French Caribbean Islands (Multigner, 2008; Multigner et al., 2006).

In humans, the acute neurotoxicity of CLD was well documented through the study of workers poisoned during the production of this pesticide at a Hopewell factory (VA, USA) in 1975. More than half of the exposed workers developed neurological symptoms, including tremors, which may involve interactions of chlordecone with neuromediators (Taylor, 1982; Taylor et al., 1978). Experimentally, oral exposure of adult mice to CLD results in dose-dependent hyperexcitability, tremors, and impaired motor coordination (Huang et al., 1981). In another study with similar experimental design, authors noticed that neurotoxicity of CLD correlated with brain and plasma concentrations of CLD (Wang et al., 1981). Although these pioneer studies highlighted the acute neurotoxicity and neurotropism of CLD, the possible neurotoxicity after chronic exposure to CLD remains to be studied.

Indeed, some pesticides have long been considered a risk factor for the development of Parkinson’s Disease (PD) (Hubble et al., 1993; Seidler et al., 1996) including through professional exposures to organochlorine insecticides in France (Dutheil et al., 2010; Elbaz et al., 2009). The concentrations of the organochlorines dieldrin and lindane were higher in PD substantia nigra (SN) compared to other neurodegenerative diseases (Corrigan et al., 2000). These results suggested that organochlorines could have a direct toxic action on the dopaminergic system and may contribute to the development of PD. A recent study showed that the presence of Lewy pathology in the brain, even after exclusion of PD cases, was associated with organochlorines, and mainly with heptachlor epoxide found at “extraordinary high” levels in milk supply as a metabolite derived from heptachlor, heavily used by pineapple growers in Hawaii after World War II (Ross et al., 2019). More recently, an association between Parkinson-like neurodegeneration and CLD was experimentally demonstrated, with dopaminergic cell losses identified in mouse midbrain cultures and in C. elegans worms (Parrales-Macias et al., 2023).

Epidemiological studies regarding a possible association of neurological diseases with long term exposure to chlordecone are still missing, although an atypical parkinsonism has been specifically described in the French West Indies (Caparros-Lefebvre et al., 2002), of still ill-defined origin (Caparros-Lefebvre et al., 2006). However, several epidemiological studies were recently reported based on a prospective longitudinal follow-up of children exposed during the gestation or post-natally in Guadeloupe. These showed that CLD was already associated with negative effects on cognitive and motor development in 7-month-old infants (Dallaire et al., 2012) and impairments in fine motor function in 18-month-old boys (Boucher et al., 2013). Further, at 7 years old, subtle hand tremors and poorer visuospatial processing was found associated with chlordecone (Desrochers-Couture et al., 2022), as well as impaired visual contrast sensitivity (Saint-Amour et al., 2020).

In this study, our aim was to set up a protocol of sub-chronic exposure of mice to study CLD neuro-accumulation and its neurological consequences. We intraperitoneally injected young adult male mice with CLD for two months and studied the population of dopaminergic neurons using a stereological immunohistochemical approach and biochemical studies. Functional consequences were assessed using behavioural tests for anxiety/depression and an innovative method of functional ultrasound imaging (fUSi), a recent neuroimaging technique that can noninvasively map changes in cerebral blood volume (CBV) at high spatiotemporal resolution in live animals (Deffieux et al., 2018; Macé et al., 2011). While fUSi has already been employed to study pharmacological effects of drugs (pharmaco-fUS) (Rabut et al., 2020; Vidal et al., 2020a, 2020b) we here highlight for the first time its potential in the field of neurotoxicology (Toxico-fUS).

## 2. Material and methods

### 2.1. Animal housing

#### 2.1.1. Housing and chlordecone injection

Animal experiments were performed at the ANSES-Lyon facility (approval number E69 387 0801), and fUSi was performed at the CERMEP-Imagerie du Vivant facility (approval number C69 383 0501), in accordance with EEC Directive 86/609/EEC and French decree No. 2013-118. The experimental protocol was authorized (APAFIS#23481-2019121811091473 v4) by the ANSES/ENVA/UPEC ethic committee (CEEA – 016). Mice were group-housed under a 12:12 light–dark lighting schedule with free access to food and water.

Six- to eight-week-old male C57BL/6JRj mice (Janvier Labs) were injected intraperitoneally with ∼1.5% dimethylsulphoxide diluted in 0.9% NaCl (100 μl each, n=22) or chlordecone (3 mg/kg in a volume of 100 μl ∼1.5% dimethylsulphoxide diluted in 0.9% NaCl each, n=34) twice a week for 8 weeks. Oral LD50 values for chlordecone range from 71 mg/kg for rabbits to 250 mg/kg for dogs (Larson et al., 1979). Body weight was measured before every injection in all animals. No formal sample calculation was performed for this exploratory study. Experimental groups are reported in **Table 1**. All analyses were done by experimenters blind to group allocation.

**Table 1.**
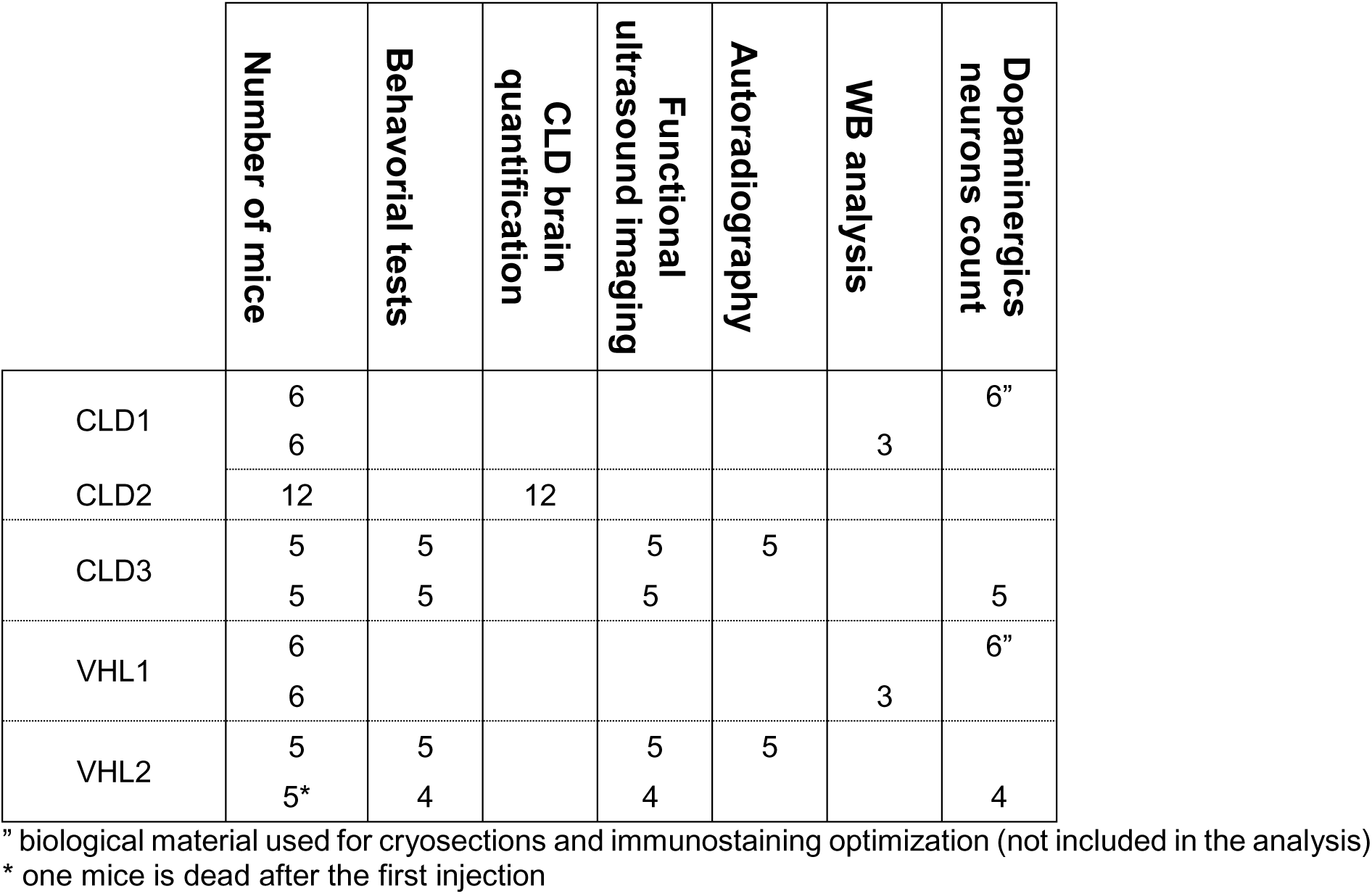
Detailed experimental groups and associated analysis. These table listed all groups of animals exposed to chlordecone (CLD1, CLD2 and CLD 3) or vehicle solution (VHL1 and VHL2). CLD1, CLD2 and VHL1 were carried out simultaneously. CLD3 and VHL2 were also performed simultaneously but in a second time (6 months apart).

#### 2.1.2. Bodyweight analysis

Evolution of the mice mass measured over time were modeled using mixed effects models as we were concerned with analysis of repeated measurements over time on the same animals (Laird and Ware, 1982).

Two models were fitted, one (model 1) for CLD1, CLD2 and VHL1 which mice were housed simultaneously and a second (model 2) for CLD3 and VHL2 were also performed simultaneously but in a second time (6 months apart). Both models included fixed effects for the groups and random effects to account for the mice variability.

Model assumptions were checked graphically by examination of the residuals, comparison of fitted and observed values and distribution of the random variables.

Analyses were performed using R software (R Development Core Team, 2009) and nlme packages.

### 2.2. Behavioural tests

Behavioral testing was performed at three time points: early (2 weeks), middle (4 weeks), and late (8 weeks) after the initiation of chlordecone (CLD, n = 10) or vehicle (VHL, n = 9) administration. All tests were conducted during the animals’ active (dark) phase, under red-light conditions.

#### 2.2.1 Anxiodepressive-like Behaviors

##### 2.2.1.1 Splash Test

The Splash test was used to evaluate depressive-like behaviors, as it reflects self-care motivation. Mice were placed individually in clean cages, and a 10% sucrose solution was sprayed on the dorsal coat. Grooming behavior was recorded for 5 minutes, including latency to initiate grooming and total grooming duration (Santarelli et al., 2003). A reduction in grooming duration is considered indicative of depressive-like behavior.

##### 2.2.1.2 Open Field Test

The Open Field test was used to assess anxiety-like behavior based on rodents’ aversion to open spaces (Prut and Belzung, 2003). Mice were placed in a 40 × 40 × 40 cm opaque chamber and allowed to explore freely for 10 minutes (Hou et al., 2021). The number of entries into the central zone was recorded, with reduced central activity interpreted as increased anxiety-like behavior.

##### 2.2.1.3 Marble Burying Test

This test measures anxiety-like and repetitive behaviors (Nicolas et al., 2006). Mice were placed individually in Plexiglas cages (40 × 34 × 19 cm) with 5 cm of clean sawdust bedding. Twenty glass marbles (1 cm diameter) were arranged in five evenly spaced rows. After 30 minutes of undisturbed activity, the number of marbles buried (≥ two-thirds covered by sawdust) was counted.

#### 2.2.2 Motor Activity

##### 2.2.2.1 Rotarod Test

Motor coordination and balance were evaluated using the Rotarod (Hou et al., 2021). Mice were placed on an accelerating rotating rod (4 to 40 rpm) divided into five compartments. Latency to fall was recorded over five trials, with a maximum of 300 seconds per trial. The average fall time was used for analysis.

##### 2.2.2.2 Pole Test

The Pole test was used to assess bradykinesia and motor agility (Fleming et al., 2004; Ogawa et al., 1985). Mice were placed head-up on top of a vertical wooden pole (50 cm tall, 1 cm diameter) and trained to descend into their home cage. During testing, five trials were performed, recording the time to orient downward (t-turn) and the total time to descend (t-total). The best trial was retained for analysis. Mice unable to turn, fell, or slipped were assigned a default time of 120 seconds.

#### 2.2.3 Learning and Memory

##### 2.2.3.1 Novel Object Recognition (NOR) test

Recognition memory was assessed using the NOR test (Leger et al., 2013). Mice were first habituated for 10 minutes in an empty opaque box (40 × 40 × 40 cm). In the training phase, they were exposed to two identical objects for 5 minutes. After a 1–2 hour retention interval, one object was replaced with a novel one, and mice were reintroduced for 5 minutes. Exploration times were recorded, and the discrimination ratio was calculated as the time spent with the novel object divided by the total exploration time.

As the same apparatus was used for the Open Field and NOR tests, activity during the habituation phase was also analyzed for anxiety-like behavior.

#### 2.2.4 Statistical analysis

Data were analyzed using a two-way repeated measures ANOVA with factors Group (CLD vs VHL) and Week (2, 4, 8) as within-subject factor. When sphericity was violated (Marble burying test), the Greenhouse-Geisser correction was applied. Post-hoc comparisons were conducted using Bonferroni correction (Splash test). For the comparison of improvement between week 2 and week 8, an independent t-test was used (Pole test).

### 2.3. Isotopic dilution liquid chromatography–tandem mass spectrometry for quantification of CLD in brain tissue

Detailed description of sample extraction procedure development, method validation and range determination were in **supplementary information**.

#### 2.3.1 Chemicals and reagents

All solutions were prepared with analytical reagent grade chemicals and ultrapure water (18.2 MΩ.cm) obtained by purifying distilled water with a Milli-Q system associated with an Elix 5 purification system (Merck Millipore, Saint-Quentin-en-Yvelines, France). Formic acid (98%) and, acetonitrile (HPLC grade), were both purchased from Fisher Scientific (Illkirch, France). CLD (98%), CDL-OH (98%) and the internal labelled standard (ILS) CLD ^13^C_10_ (98%) standards were purchased from Cluzeau Info Labo (Sainte-Foy-La-Grande, France). Working solutions were prepared by dilution of the commercial standards in acetonitrile before use, for calibration standards and fortification of quality controls and samples for method validation. All solutions were stored at 5 ± 3°C.

Dispersive Solid Phase Extraction (dSPE) step was performed using a mixture of 150 mg of C18, 150 mg of PSA and 900 mg of MgSO_4_ (Fatty Sample EN, Agilent, Les Ulis, France) and a mixture of 150 mg of PSA and 900mg of MgSO_4_ (Fruits and Vegetables AOAC Agilent, Les Ulis France). Syringe filters 0.22 µm Nylon filter (Xtra PTFE 20/25, 0.020 µm) were purchased from Macherey Nagel (Hoerdt, France).

#### 2.3.2 Equipment

LC-MS/MS measurements were performed using a 1200 series HPLC binary pump system (Agilent Technologies, Les Ulis, France) coupled to an API 5500 QTRAP hybrid tandem mass spectrometer (AB Sciex, Les Ulis, France). An Aqua C18 column (150 mm x 2.0 mm i.d., 3 µm particle size) equipped with an Aqua C18 guard column (4.0 mm x 3.0 mm i.d., 3 µm particle size) were used for chromatographic separation.

***Sample preparation and extraction.*** After grounding with the same portion of ultrapure water with micro-beads using a sample homogenizer (Bertin Technologies), extraction was performed using an Eppendorf centrifuge 5810 (Hamburg, Germany) and a Genie 2 vortex supplied by Scientific Industries (Bohemia, NY, United States). After weighing the grounded brain region in a 50 mL centrifuge tube, different test portions of ACN for extraction were added. The sample was stirred for 1 min and CLD- ^13^C solution of 500 ng.ml^-1^ was added and kept in contact for 2 hours. For dSPE experiments, 6 mL of the supernatant was collected and deposited into a 15 mL centrifuge tube. With or without dSPE, the samples were vortexed for 1.0 min and then centrifuged at 4000 rpm for 5.0 mins. Lastly, the samples were filtered on a 0.20 µm PTFE Chromafil filter and deposited in a 2.0 mL amber vial for analysis.

#### 2.3.3 LC-MS/MS analysis

LC-MS/MS analysis were adapted from Lavison et al,. 2021. The volume injected was increased from 5 to 10 μL, the column kept at 40 °C. The flow rate was set at 0.4mL/min, and the gradient adapted to limit analysis time (from 11 min to 6 min for separation). The elution was performed with mobile phases containing ultrapure water with 0.1% (v/v) formic acid (A) and acetonitrile with 0.1% (v/v) formic acid (B). The gradient starts at 75% B and remains constant for 4.5 minutes. The mobile phase B increases to 95% in 1 minute and remains constant for 1 minute before decreasing in 1 minute to 75% B followed by 3.5 minutes for equilibration of the column. Analyst 1.51 Software (AB Sciex) was used for system control and data acquisition and processing. In each sample batch, quality controls were included, with criteria recommended by SANTE guidelines (SANTE/2020/12830. Rev 2 – 14 February 2023, 2023). Bracketing calibration (5 levels) was performed, and the calibration model was accepted through residual deviation below 20%. Identification requirement on both retention times ion ratio deviations (± 30%) was systematically examined. Results were validated only if all criteria were within the acceptable limits. Analyses were performed under the international standard ISO/IEC 17025:2017.

#### 2.3.4 Statistical analysis of CLD concentrations

Normal distribution was assessed with Shapiro-Wilk tests and followed by ANOVA tests. The statistical analysis was performed with Studio software, v 1.2 (R Development Core Team, 2016).

### 2.4. Functional ultrasound imaging

#### 2.4.1 Acquisitions

For fUSi, mice were anesthetized with ketamine/medetomidine (70 mg/kg i.p./0.5 mg/kg i.p.) and scanned (according to randomization table) with a device dedicated to small animal ultrasound neuroimaging (Iconeus, Paris, France) while maintained in a stereotactic frame. Hair on the skull was removed (by shaving and by applying depilatory cream), and contact with the probe was ensured with ultrasound gel. Doppler vascular images were obtained using the Ultrafast Compound Doppler Imaging technique. Each frame was a Compound Plane Wave frame (Montaldo et al., 2009) resulting from the coherent summation of backscattered echoes obtained after successive tilted plane waves emissions. A stack of hundreds of such compounded frames was acquired with a high frame rate. Each transcranial Doppler image was obtained from 200 compounded frames (Montaldo et al., 2009) of Doppler vascular images (Bercoff et al., 2011) acquired at 500 Hz frame rate. A fast scan with successive images taken on several coronal planes was performed for positioning the probe in the planes of interest.

A first acquisition was performed under visual stimulation. The probe was placed at -3 mm from bregma. Images were acquired every 400 ms during a total of 405 seconds. Visual stimuli were delivered using a screen in front of the mice. Stimulation runs consisted of black and white flickering on the screen for stimulation (at random frequencies ranging from 0.5 to 15 Hz) and continuous black screen for rest. The stimulation pattern consisted in 30 s of initial rest followed by runs of 30 s of flicker and 45 s of rest, repeated five times. This was performed in the dark after waiting for at least 15 min, to ensure that the mice were dark-adapted.

A second acquisition was performed during whisker stimulation. The probe was placed at -1.5 mm from bregma. Again, images were acquired every 400 ms during a total of 405 seconds. The stimuli were manually delivered using a Q-tip to brush the left whiskers, with a block pattern identical to the visual stimulation (30s of initial rest followed by runs of 30 s of stimulation and 45 s of rest, repeated five times).

Finally, mice underwent a pharmacological challenge during a third acquisition, in order to follow the hemodynamic changes induced by a stimulation of the dopaminergic system. The probe continuously moved between bregma +1 mm and bregma -3 mm, at a rate of 1 image every 2.8s for each plane. The total duration of the acquisition was 35 minutes, with 5 minutes of baseline followed by an intraperitoneal injection of apomorphine (1 mg/kg) and 30 minutes of post-injection period.

The total duration of anesthesia for fUSi was in the range of 45-75 min, and awakening was facilitated by the injection of atipamezole (4 mg/kg s.c.).

#### 2.4.2 In vivo imaging analysis

Before data processing, all acquisitions were automatically registered together in Matlab, as previously described (Vidal et al., 2021). After the registration, the corresponding two-dimensional slices of interest (+1 mm, -1.5 and -3 mm from bregma) were extracted from the Allen atlas and manually co-registered on the template using ITK-Snap software. The CBV changes were extracted from the regions of interest after temporal frequency filtering (highpass and lowpass filters with cutoffs frequencies of 0.01 and 0.15 Hz for visual and whisker stimulations, and only a lowpass filter with a cutoff of 0.15 Hz for pharmacological stimulation).

For visual and whisker stimulation, the mean CBV changes during all stimulations were calculated and the five time-courses were averaged to estimate a global CBV time-course during the sensory stimulation for each acquisition, before averaging all individuals for each group. For all stimulations, the global mean CBV changes occurring during the whole stimulation period (mean of the five blocks or whole post-injection period for the pharmacological stimulation) as compared to the baseline were calculated for each fUS acquisition. For the final statistical analysis, two-way ANOVAs were used to compare mean CBV changes in the different regions of interest between the VHL and CLD groups, followed by multiple comparisons with an false discovery rate (FDR) threshold of 0.1.

### 2.5. In vitro autoradiography

10 animals enrolled in the fUSi protocol were used for autoradiography (n=5 each). Upon sacrifice under ketamine/xylazine anesthesia, brains were extracted, snap-frozen in isopentane, and stored at -80°C. 20µm-thick brain sections were cryostat-cut at the level of the striatum and substantia nigra and stored at -80°C on microscopy slides. Radiosynthesis of [^11^C]PE2I and [^11^C]raclopride was performed as per standard procedure at the CERMEP-Imagerie du Vivant. Brain sections were incubated with radiotracers (1µCi/mL) in PBS for 20 minutes (to measure total binding), and adjacent sections were incubated with the same amount of radiotracer together with a excess of non-radiolabelled competitor: 100 µM GBR12909 for [^11^C]PE2I, 10 µM haloperidol for [^11^C]raclopride (to measure non-specific binding). All slides underwent two washing steps of 5 min, before being dried under cool air and exposed to a sensitive phosphor-imaging plate (BAS-IP MS 2025, Fujifilm) for two hours. The distribution of radioactivity was then digitized with a bio-imaging analyser system (BAS-5000, Fujifilm).

Quantification was performed with dedicated software (Multi Gauge, Fujifilm). Total and non-displaceable binding were extracted from bilateral ellipsoid regions of interest manually placed in the center of the striatum or substantia nigra, and identically reported on the corresponding section with competitors. Five brain sections were averaged for each animal and each radiotracer. A two-way ANOVA was used to compare the binding in the striatum and substantia nigra between the VHL and CLD groups.

### 2.6. Western Blot

Protein extractions were obtained from dissected brain areas homogenized at 10% in High Salt (HS) Buffer (50 mM Tris–HCl pH=7.5, 750 mM NaCl, 5 mM EDTA, 1 mM DTT, 1% phosphatase and protease inhibitor cocktail (ThermoFisher #1861281). Brain homogenates were mixed with TD4215 denaturing buffer (final concentration: 4% SDS, 2% β mercaptoethanol, 192 mM glycine, 25 mM Tris, 5% sucrose). Then, after heat denaturation (5 min at 100°C), samples were loaded in acrylamide TGX Stain Free Precast gels (5678044 Biorad, Marnes-La-Coquette, France) gels with 20 µg proteins per lane. Proteins concentrations were determined with DC protein Assay (Biorad, Marnes-La-Coquette, France). After migration step, Stain Free gels were activated for 1 min on Chemidoc MP apparatus (BioRad, Marnes-La-Coquette, France) and image analysis was done using ImageLab software (BioRad) to control the protein loads per lane. Then, proteins were blotted onto PVDF membranes (Immobilon-P, Sigma, Saint-Quentin Fallavier, France) and were cross-linked to the membranes using 4% paraformaldehyde (40877-29-MMF, Micro Microtech) and 0.01% glutaraldehyde (340855-25ML, Sigma Aldrich) for 30 minutes. After, unspecific binding sites were saturated by adding 5% milk for 1 h. Five different protein detections were carried out with specific primary antibodies such as Tyrosine Hydroxylase (TH, Abcam #Ab 112, at 1/5000), Dopamine Transporter (DAT, Santa Cruz Biotechnology #Sc-32258 at 1/500), Vesicular Monoamine Transporter (VMAT2, Santa Cruz Biotechnology #Sc-374079 at 1/500), Glial fibrillary acidic protein (GFAP) for astrocytes (DAKO, #Z0334, 1/5000) and ionized calcium-binding adapter molecule 1 (IBA1) for microglia (WAKO, #01620001, 1/500). Antibodies were incubated overnight at +4°C. After washings, membranes were then incubated with secondary mabs at 1/20 000 raised against rabbit (for TH, GFAP or IBA1), rat (for DAT) or mouse (for VMAT2) (Southern Biotech, #4010-05, #3030-05 and #1010-05 for rabbit, rat and mouse respectively) and HRP-coupled for 1 h at room temperature (RT). HRP was then visualized with chemiluminescent substrate (Supersignal WestDura, 10445345, Fisher, Illkirch, France) and analyzed using the ChemiDoc system and ImageLab software (Bio-Rad). TH, DAT AND VMAT2 proteins detection have been carried out in the same membrane. Due to their difference in molecular mass TH and DAT were detected in sequence: TH followed by DAT. Removal of primary and secondary antibodies from the membrane was performed in a stripping solution (60 mM Tris–HCl pH=6.5, 0.7% β mercaptoethanol, 2% SDS) applied 30 minutes at 50°C. GFAP and IBA1 protein detection have been detected in two independents membranes. A two-way ANOVA was used to compare the different protein expression between the VHL and CLD groups.

### 2.7. Unbiased estimation of dopaminergic neuron population

#### 2.7.1 Immunostaining

An unbiased stereological estimation method (West et al 1991), adapted from previously published studies (Ip et al., 2017) or tools (Mironov, 2020), was used to assess the level of neurodegeneration within the substantia nigra pars compacta (SNpc). After euthanasia, mice were perfused transcardially with 15 mL of phosphate-buffered saline (PBS) using an infusion pump, followed by 50 mL of 4% paraformaldehyde (PFA) in PBS. The brain was removed and immersed one night at +4°C in 4% PFA solution. After PBS washes, the mouse brain was dissected in the coronal plane at the region comprises between -2 mm and -4 mm from Bregma. The central macro-section of the brain, including the SNpc, was immersed in 30% sucrose/PBS solution for cryo-protection and incubated up to 2 days at 4 °C. It was then placed into a cryomold filled with optimal cutting temperature (OCT) compound and frozen on dedicated area in cryostat apparatus (Leica CM 1860 UV). At this step, molded pieces could be conserved for several months at -80°C. 30-µm serially cryo-sections were directly immerged in a PBS solution in a 12-wells culture plate.

Immunohistochemical Tyrosine Hydroxylase (TH) staining was carried out on free-floating sections, starting with an incubation for 5 minutes in a 3% H2O2 solution to block endogenous peroxidases. This step was followed by a 30-minute incubation in 10% normal goat serum (NGS)/2% bovine serum albumin (BSA)/0.5% Triton100 detergent in PBS. The primary rabbit anti-mouse TH (1:2,000 dilution) (Abcam, #Ab112) antibody was diluted in 2% NGS/2% BSA/0.5% detergent in PBS and incubated overnight at 4 °C. After PBS washes, horseradish peroxidase (HRP) coupled secondary antibody against rabbit immunoglobulin (1:1000 dilution) (Clinisciences, #4010-05) was incubated for 2 h at room temperature. Final staining was obtained through a 3-minute incubation in a solution of 3,3-diaminobenzidine-tetrahydrochloride (DAB) and peroxidase substract (Vector Laboratories, #SK 4105, ImmPACT® DAB Substrate Kit, Peroxidase). Sections were mounted with non-aqueous mounting medium (ThermoScientific, #4112).

#### 2.7.2 Microscopy and quantification

Images were acquired using a confocal scanner (Yokogawa CQ1). After framing the fields containing the SNpc in X and Y, the center of each slice was determined using Z focus. From the latter, five 2 µm-thick optical sections were obtained above and below this central section, to yield a 22 µm-thick captured volume. The following steps were carried out using image J software and the toolkit developed by Mironov (Mironov, 2020). Briefly, the 11 optical slices were flattened using the maximum intensity method, thereby representing all the labeled neurons in the volume. The counting frame was then applied to the resulting 2D image with a random start anchor. Neurons present in the frame were counted manually. Groups were compared with a non-parametric Mann-Whitney test.

### 2.8. Immunohistochemical GFAP and IBA1 staining on cryosections

Immunolabeling of GFAP and IBA1 proteins was performed on cryosections prepared during the stereology approach, plus additional sections of other neuroanatomical areas were produced. Labeling was implemented in the same way as TH protein labeling in the stereological analysis. The primary antibodies used were (Abcam #ab7260X at 1/2000) and (WAKO #W1W016-20001Y at 1/2000) for GFAP and IBA1 respectively, and the secondary antibody was (Southern Biotech #4010-05 at 1/500) for both.

## 3. Results

All animals in both groups received two intraperitoneal injections per week. The control group (VHL) received an injection of ≈ 1.5% DMSO diluted in saline solution, and the chlordecone group (CLD) received an identical diluting solution with a chlordecone amount of 3mg/kg. In order to respect this dosage the animals were weighed at each injection. This method enabled us not only to maintain a constant dose of chlordecone, but also to monitor the general condition of the animals. Among the 56 mice of this experiment, only one mice from VHL group died between the first and second injection, the necropsy did not revealed damage associated with the protocol. All other mice completed the 8-weeks protocol, without adverse effect associated with repeated ip injections. No sign of suffering or aggression was noted.

The average weights of each groups over the eight weeks of exposure were presented in the figure 1(**Figure 1**). Evolution of the mice mass measured over time were modeled using mixed effects models. We compared in two models two groups of mice housed simultaneously: CLD1, CLD2 and VHL1 in model 1 and CLD3 and VHL2 in model 2. We compared the average mass of each group at the start of the experiment (Intercept) and the progression of mass over time (Time). The full results of the statistical analyses are shown in the supplementary figures **(Supplementary Figure 3, 4 and 5)**. In model 1, the mean mass at the origin (Intercept) for the VHL1 group is 28.05, there is no significant difference with CLD2 (+0.11 - p-value = 0.8543), on the other hand the mean mass of CLD1 is significantly lower at the origin (−1. 42 - p-value = 0.0310). In the growth analysis, there was no significant difference in growth between the three groups of animals (Time CLD1: +0.008 - p-value = 0.3062) (Time CLD2: - 0.011 - p-value = 0.1658) **(Supplementary Figure 3)**. Still in model 1, but taking CLD1 as a reference, the mean mass at origin (Intercept) for the CLD1 group is 26.63, with a significant difference in mass also observed with CLD2 (+1.53 - p-value = 0.0203). Mass gain was significantly slower in CLD2 (Time CLD2: −0.019 - p-value = 0.0162) than in CLD1 **(Supplementary Figure 4)**. In Model 2, comparing CLD3 and VHL1 groups, the mean mass at origin (Intercept) for the VHL2 group is 20.96, no significant differences were observed either at origin or in mass gain over time **(Supplementary Figure 5)**.

**Figure 1.**
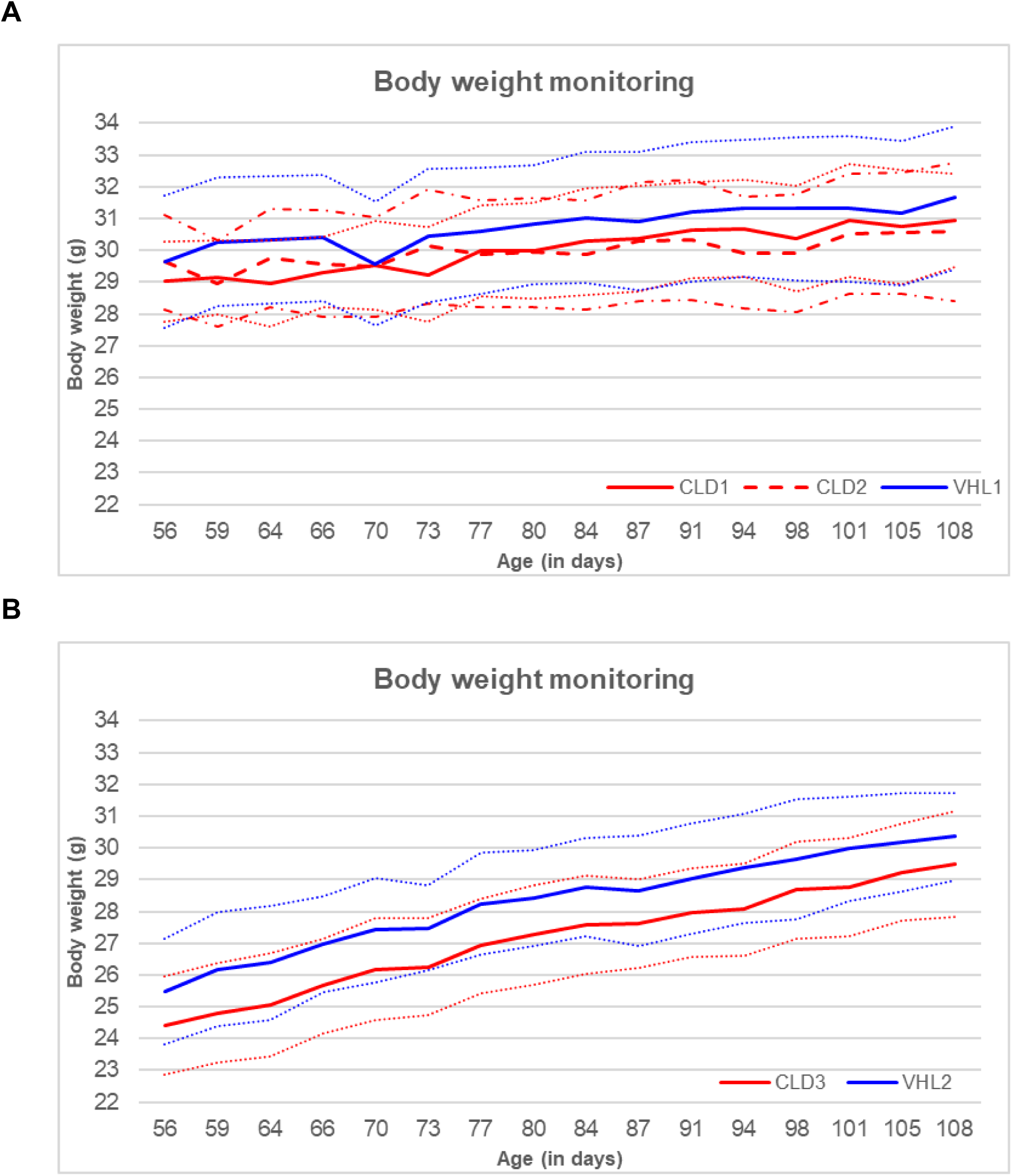
Monitoring of body weight of mice exposed to chlordecone: CLD1 (n=12), CLD2 (n=12) and VHL1 (n=12) for the first period of experimentation **(A)** and CLD3 (n=10) and VHL2 (n=9) mice for the second period carried out 6 months later **(B)**. The exposure protocol consisted of intraperitoneal injections twice a week, with the injected dose calculated from individual body weight. Solid lines correspond to the mean values for both groups, and dot lines show the standard deviation.

The distribution and quantification of chlordecone in mice brain tissues were determined by isotopic dilution liquid chromatography–tandem mass spectrometry. The complete analytical protocol and method validation are provided as supplementary information. The brains of 12 CLD mice were dissected into 10 different regions: olfactive bulb (OB), left cortex (CTXL), right cortex (CTXR), striatum (STR), hippocampus (HIP), thalamus (THAL), hypothalamus (HYPO), mesencephalon (MES), cerebellum (CRB) and brainstem (BS). The resulting 120 exposure samples were analyzed with the above-mentioned protocol and are presented in **figure 2**. All brain samples were free of CLD-OH (below the quantification limit) as expected, since mice cannot metabolize CLD into CLD-OH. For CLD, all brain regions were contaminated between 5.94 and 14.96 mg/kg, in the validated range of concentration measurement of the method, without significant difference between regions (p=0.13).

**Figure 2.**
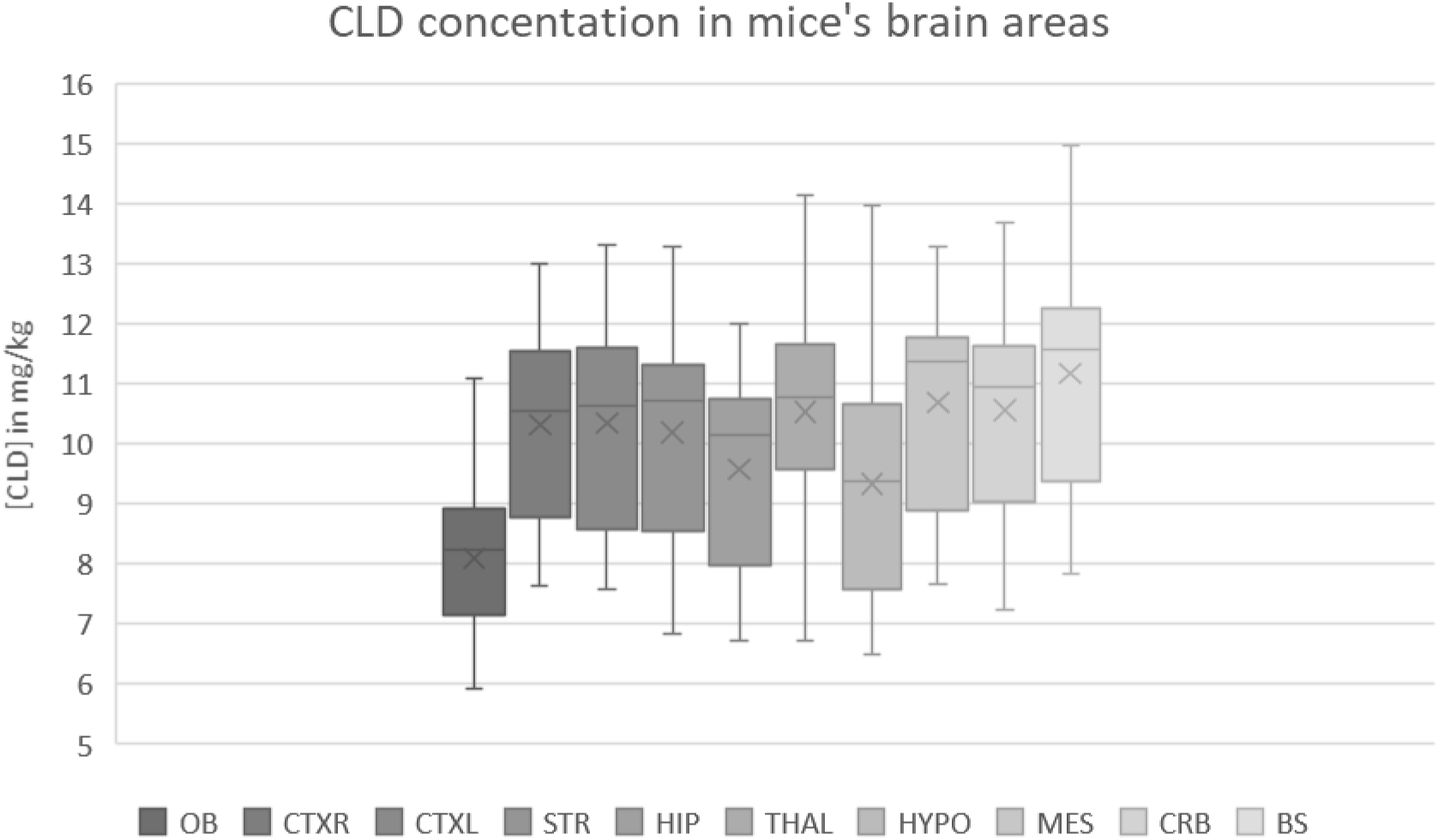
Chlordecone (CLD) concentrations (in mg/kg) in different areas of the brain. (n=12 mice): olfactive bulb (OB), left cortex (CTXL), right cortex (CTXR), hypothalamus (HYPO), striatum (STR), cerebellum (CRB), brainstem (BS), mesencephalon (MES), hippocampus (HIP) and thalamus (THAL). Finally, 119 of the 120 planned samples have been analyzed because the olfactive bulbs of one mice were destructed during the dissection step. In this graphic, the central lane is the median and the cross corresponds to the mean value. The top side of the box represents the third quartile and the bottom side the first. The upper bar displays the maximum value of the series and the lower bar the minimum value.

No significant differences were observed between the CLD and VHL groups in measures of depressive- or anxiety-like behavior. Specifically, grooming duration in the Splash test was similar between groups (Supplementary Figure S2A). In the Open Field test, both groups made a comparable number of entries into the center zone (Supplementary Figure S2B). Likewise, marble burying behavior did not significantly differ between groups (Supplementary Figure S2C).

Short-term memory, as assessed by the Novel Object Recognition test, was also unaffected by CLD exposure, with no significant group differences in discrimination ratios (Supplementary Figure S2D). In terms of motor performance, the Rotarod test showed no significant differences in latency to fall between groups at any time point (Supplementary Figure S2E).

Similarly, the Pole test revealed no differences in performance between groups when comparing individual time points. However, longitudinal analysis of test performance revealed a significant group difference in agility improvement: between weeks 2 and 8, VHL mice exhibited a marked reduction in descent time, whereas CLD-exposed mice did not (p = 0.031; **Figure 3**). These findings suggest that although chronic CLD exposure does not overtly impair motor coordination or learning, it may hinder the refinement of agility that typically improves with task repetition.

**Figure 3.**
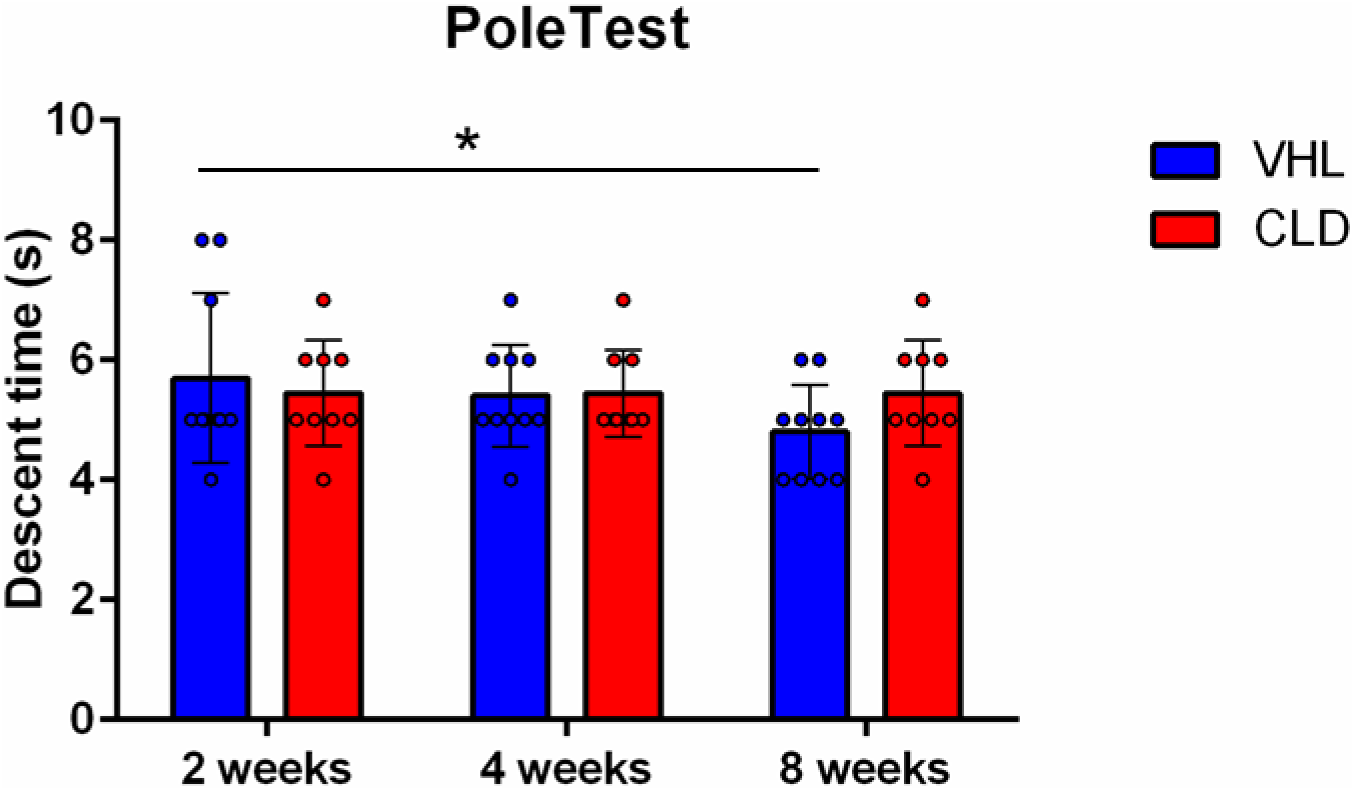
Improvement in performance is impaired in the Pole test following chronic chlordecone exposure. Individual points show the best performance (shortest time in seconds) out of five trials in the Pole test at weeks 2, 4, and 8 of exposure for vehicle-treated (VHL, n = 9) and chlordecone-treated (CLD, n = 10) mice. The test measured the total time taken to turn downward and descend from the pole. Data are presented as mean ± SD. *p < 0.05.

The functional consequences of this chronic chlordecone exposure were assessed using functional ultrasound imaging (fUSi), a neuroimaging technique that can non-invasively map the changes in cerebral blood volume (CBV) in mice. Three distinct stimulation protocols were used in order to probe the functionality of different pathways, namely the visual pathway (**Figures 4A-C**), the somatosensory pathway (**Figures 4D-F**), and the dopaminergic pathway (**Figures 4G-I**).

**Figure 4.**
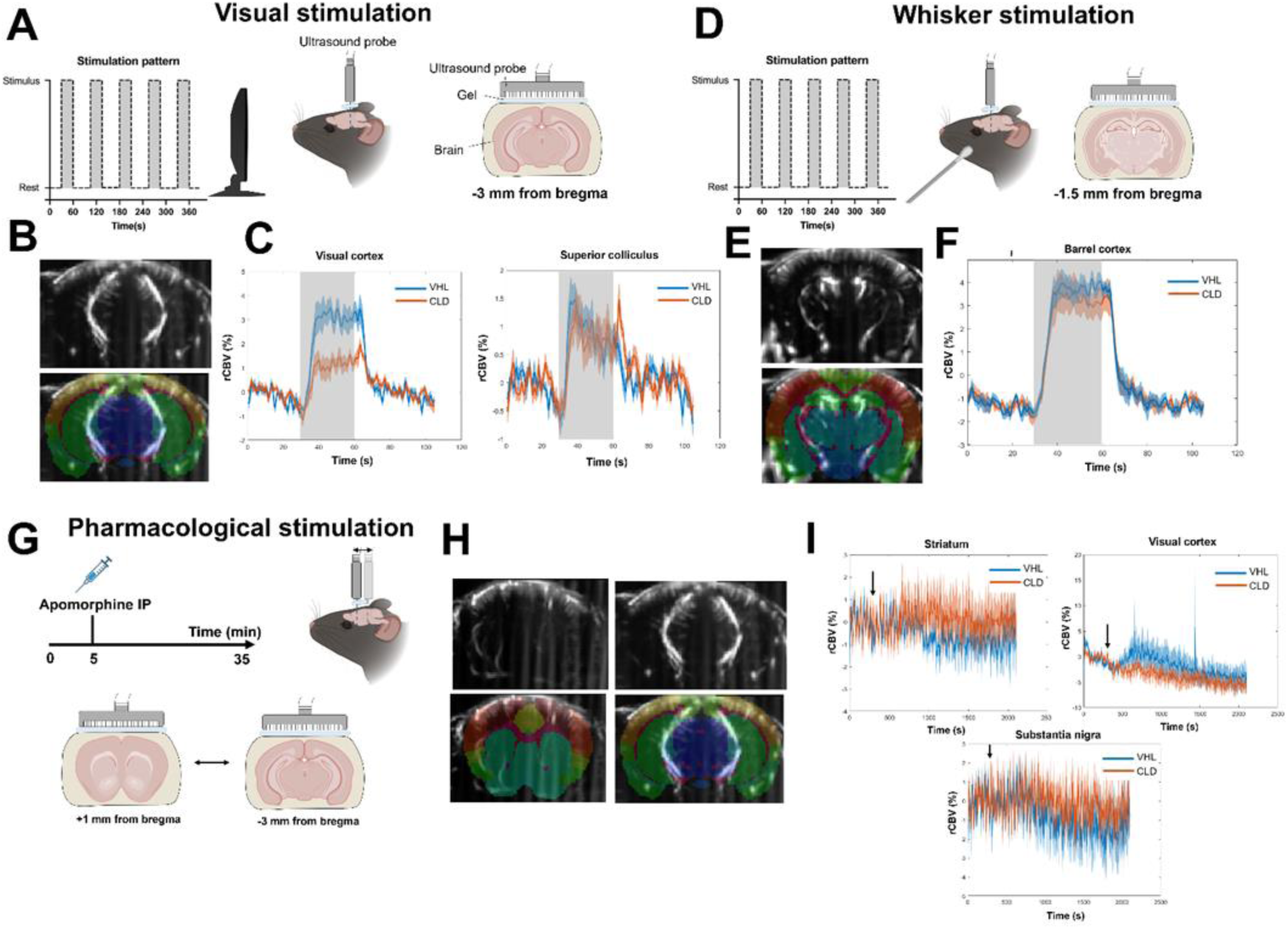
Changes in brain activity after exposure to chlordecone (CLD, n=10) vs vehicle (VHL, n=9) studied by functional ultrasound imaging (fUS). (**A**) fUS acquisition during visual stimulation. (**B**) Example of fUS image at -3 mm from bregma during visual stimulation (top) and overlay of the anatomical atlas (bottom). (**C**) Mean time curves of CBV changes during all five stimulation periods in the anterolateral visual cortex (left) and gray layer of the superior colliculus (right). (**D**) fUS acquisition during whisker stimulation. (**E**) Example of fUS image at -1.5 mm from bregma during whisker stimulation (top) and overlay of the anatomical atlas (bottom). (**F**) Mean time curves of CBV changes during all five stimulation periods in the barrel field of the primary somatosensory cortex. (**F**) fUS acquisition during pharmacological stimulation. (**H**) Example of fUS images at +1 (left) and -3 mm (right) from bregma during pharmacological stimulation (top) and overlay of the anatomical atlas (bottom). (**I**) Mean time curves of CBV changes during the whole acquisition in the striatum (top left), the anterolateral visual cortex (top right) and the substantia nigra pars compacta (bottom).

To assess the visual pathway, stimuli were applied using a screen during a first fUSi acquisition at -3 mm of bregma (**Figure 4A**), in a block design that alternated between rest periods (black screen) and stimulation periods (light flickering). All individual images were registered in the same space before co-registration with the corresponding coronal slice from the Allen atlas (**Figure 4B**), enabling to extract the time-courses of relative CBV changes in different regions of interest. The mean temporal response, averaged between the five stimulation blocks, is shown in **Figure 4C** for some typical regions of the visual pathway (left: anterolateral area of the visual cortex, layer 4; right: superficial gray layer of the superior colliculus) to illustrate the specific impact of chlordecone treatment in visual cortical areas as compared to subcortical regions. All individuals mean amplitude of response in all regions of interest can be found in **Supplementary Figure 6**. A significant decrease in the mean response amplitude (in the CLD vs VHL mice) was found in several visual cortical areas, such as the anterolateral area (−0.95%, -1.64% and -0.78%; q=0.0198, q<0.0001 and q=0.0956 in layers 2-3, layer 4 and layer 5, respectively) and the posteromedial area (−0.98%, q=0.0174 in layer 4). Some areas of the posterior auditory cortex also displayed some functional hyperemia during visual stimulation, with a response that was significantly lower in the CLD group (−0.81% and -0.98%, q=0.0835 and q=0.0174 in layers 2-3 and layer 4, respectively). All other visual cortical areas – namely all layers of the primary visual cortex, and layers 1 and 6 of the anterolateral/posteromedial visual cortex – also tended to display a weaker response to visual stimuli in the CLD group, but the difference were not statistically significant (q>0.1). The visual-related areas of the superior colliculus (optic layer, superficial gray layer and zonal layer) also displayed an increase of CBV during visual stimulation, as expected, but the mean amplitudes were not statistically different between groups (q>0.1), although a late peak was observed in the time courses of CBV changes at the end of the stimulation for the CLD group only (**Figure 4C, right**).

To assess the somatosensory pathway, stimuli were applied manually using a Q-tip to brush the left whiskers, during a second fUSi acquisition at -1.5 mm of bregma (**Figure 4D**), in a block design that alternated between rest periods (no brushing) and stimulation periods. As before, all individual images were registered in the same space before co-registration with the corresponding coronal slice from the Allen atlas (**Figure 4E**) to extract the time-courses of relative CBV changes in different regions of interest. A typical example of the mean temporal response, averaged between the five stimulation blocks, is shown in **Figure 4F** (primary somatosensory area, layer 1 of the barrel field). As expected, there was an increase of CBV in different layers of the barrel field, but also in the trunk field and the dorsal part of the retrosplenial area, to a lesser extent. In contrast with the visual stimulation, the somatosensory stimulation yielded no significant change in amplitude in any of those regions between the CLD and VHL groups (as illustrated in **Figure 4F** for the barrel field).

Finally, a pharmacological stimulation of dopaminergic receptors was performed in the same mice during a third fUSi acquisition using an intraperitoneal injection of apomorphine at 1 mg/kg (**Figure 4G**). During this scan, the probe was continuously displaced between -3 mm of bregma and +1 mm of bregma to assess the CBV changes in the striatum as well as the substantia nigra. Again, all images for each plane were registered in the same space before co-registration with the corresponding coronal slice from the Allen atlas (**Figure 4H**) to extract the time-courses of relative CBV changes in anatomical regions. The temporal response over the 5 minutes of baseline and 30 minutes following apomorphine injection is shown in **Figure 4I** for different regions of interest, namely the striatum, the substantia nigra pars compacta and the visual cortex (anterolateral visual area, layer 4, as in **Figure 4C**). 5 individual acquisitions (3 VHL, 2 CLD) were excluded due to technical problems during the pharmacological challenge. No significant difference of global post-injection CBV changes was found between the CLD and VHL groups following apomorphine stimulation, despite the average time courses tended to differ (for instance, a slight decrease of CBV was observed in the striatum and the substantia nigra pars compacta for the vehicle group only; conversely, a slight increase of CBV was found in the visual cortex, which was specific of the vehicle group – **Figure 4I**).

To examine whether chlordecone induced degeneration of dopaminergic neurons in the SN pars compacta (SNpc), five animals from CLD group and four animals from VHL group were sacrificed 1 week after the last injection to perform TH immunostainings. When the numbers of TH-positive neurons were counted in the SNpc area, CLD mice showed a non-significant decrease (−15%) in the TH-positive dopaminergic neuronal population compared to VHL mice (**Figure 5**). Groups were compared with a non-parametric Mann-Whitney test.

**Figure 5.**
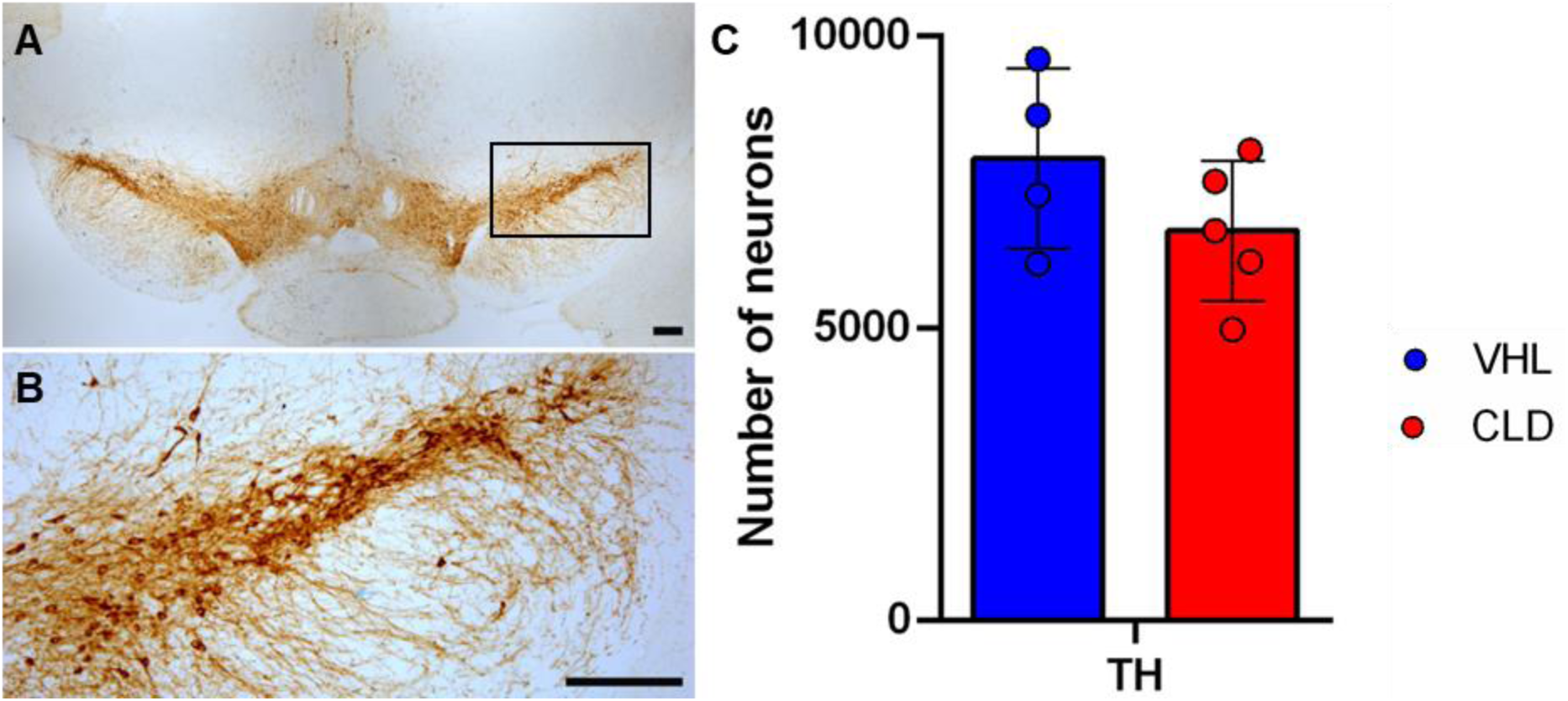
Stereological estimation of the number of dopaminergic neurons in mice exposed to chlordecone. (CLD, n=5) or vehicle (VHL, n=4). Unbiased stereology was performed to count the number of dopaminergic neurons in the SNpc of the mice after 8 weeks of exposure. TH immunoreactive cells in the midbrain (mice from VHL group), stained brown using DAB at microscope objectives x5 (**A**) and x10 (**B**), scale bar represent 200µm for both pictures. The box in the panel A represent the field observed at x10 in the following panel. The stained sections were observed with a light microscope BX51 (Olympus) coupled to INFINITY3-6UR camera (Lumenera) and Lumenera software (Infinity analyze version 6.5.5) was used. (**C**) DAB-stained cell counts for each group. Groups were compared with a non-parametric Mann-Whitney test.

Further quantitative assessment of the dopaminergic system included i) the determination of the protein levels of Tyrosine hydroxylase (TH), dopamine transporter DAT (DAT) and Vesicular monoamine transporter 2 (VMAT2) in the striatum by immunoblot assay (**Figure 6A-B**), and ii) the measurement of the densities of dopamine transporter and D2-receptors in the striatum and in the substantia nigra (SN) by in vitro autoradiography (**Figure 6C-D**). All these protein parameters remained unchanged except VMAT2 which decreased in the striatum of the three exposed mice analysed. We also studied neuroinflammatory responses using markers associated with astrocytes (GFAP) and microglia (Iba1) either by western blot or IHC in different brain areas such as the striatum, olfactory bulbs, midbrain and cortex. No significant variations were observed between groups (data not shown).

**Figure 6.**
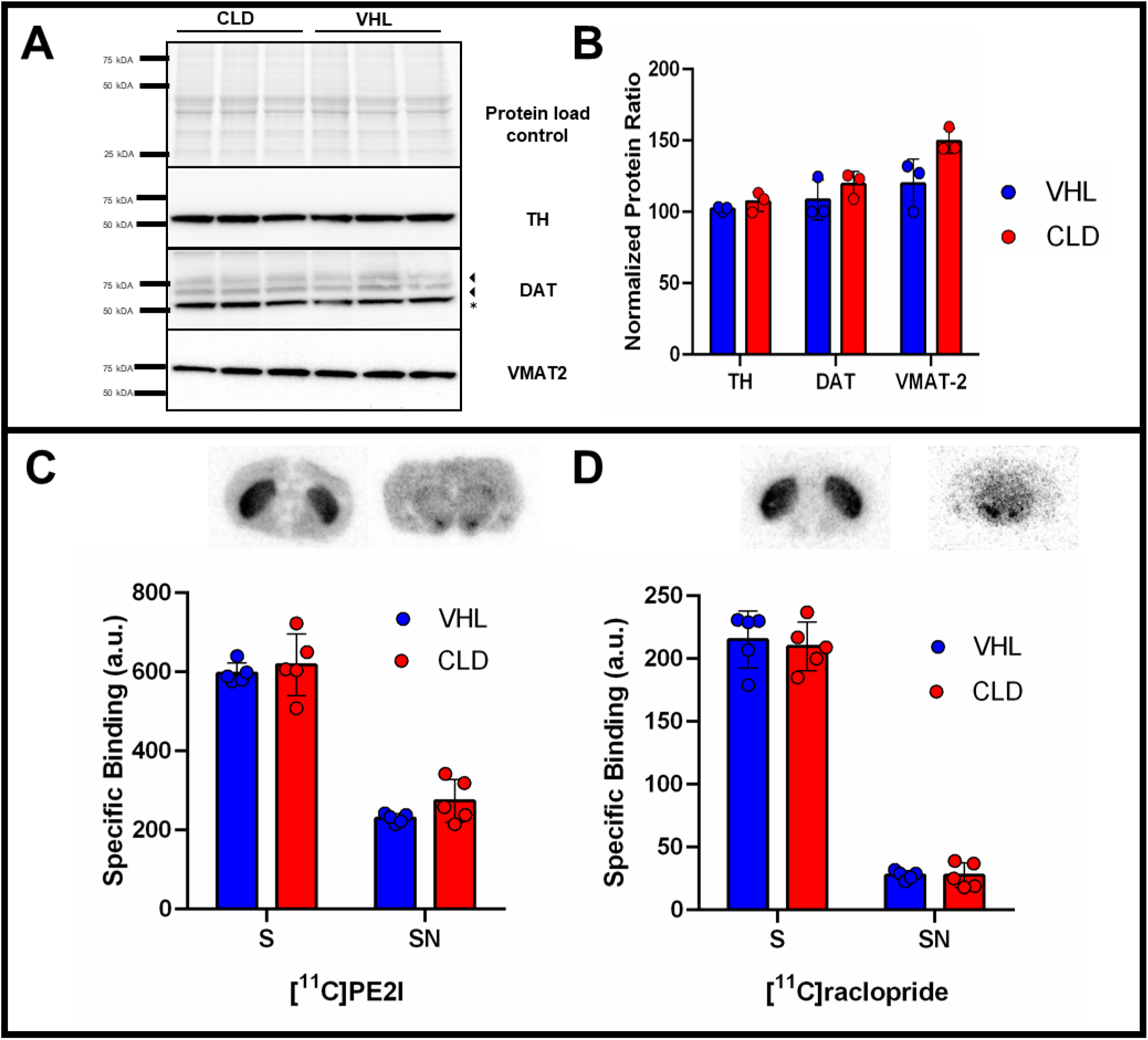
Impact of chlordecone exposure on the dopaminergic system. (**A**) Western Blot for tyrosine hydroxylase (TH), dopamine transporter DAT (DAT), vesicular monoamine transporter 2 (VMAT2) in the striatum (n=3 per group), and (**B**) associated quantification. (3 : DAT bands usually visualized, *: bands associated to TH signal previously immuno-labelled in this membrane as presented below) *In vitro* autoradiography using radiotracers binding to the dopamine transporter, [^11^C]PE2I (**C**), and D2-receptors, [^11^C]raclopride (**D**), performed at the level of the striatum (S) and substantia nigra (SN). Quantification was carried out bilaterally using 5 sections per animal and 5 animals per group, after non-specific binding assessment using GBR12909 (for [^11^C]PE2I) and haloperidol (for [^11^C]raclopride). (black dots, VHL = vehicle; white dots, CLD = chlordecone)

## 4. Discussion

Using a sub-chronic CLD exposure model in C57Bl/6 mice, we observed a discrete deleterious effect on the population of dopaminergic neurons in the *substantia nigra pars compacta* (SNpc). Most importantly, by ultrasound functional imaging in live animals, we found a significant functional impairment of several cortical areas associated with vision. These experimental observations suggest that CLD could be associated with early alterations characterizing Parkinson’s disease or other neurological deleterious effects. This is echoing with a wealth of recent epidemiological data showing neurological alterations in children pre- or/and postnatally exposed to CLD in the French West Indies, where this chemical impregnates the general population.

Whereas the well-known acute neurotoxicity of CLD for humans has been reproduced experimentally in rodents, the general aim of our study was to initiate investigations on the neurological consequences of longer CLD exposures in mice. For this, we chose a 3 mg/kg dose administrated twice per week by intraperitoneal injections to C57Bl/6 male mice during two months. Experiments carried out in mice during the 80’s, focusing on acute neurotoxicity, used unique or repeated CLD doses, comprised between 10 and 50 mg/kg/day, administered by oral route (Huang et al., 1981; Fujimori et al., 1982; Benet et al., 1985). This may at least partly explain our different observations about CLD distribution in the brain after 2-month exposure by intraperitoneal route. Although this first confirmed the high neurotropism of CLD, we indeed rather observed an even distribution of CLD in the nine brain areas examined, with concentrations comprised between 6 and 15 µg/g of brain tissue. Instead, Fujimori et al 1982 and Benet et al 1985 found a significantly higher level of chlordecone in the striatum and medulla/pons than in the other brain regions studied after acute oral exposure. Regarding the body weight and behavioral monitoring, our protocol was well tolerated by the mice, so that longer exposures could thus be considered in the future. In another study of orally exposed mice (Wang et al., 1981), the cerebral threshold level of CLD associated with loss of motor coordination was estimated at 10-65 µg/g. Whereas CLD concentrations of 6 - 15 µg/g were observed in our conditions, no overt clinical manifestations were observed, and only the pole test was able to reveal a reduced improvement of the capacity of the CLD-exposed mice through the repeated tests during the 2-month protocol.

Our study has several limitations related to the chosen protocol. Firstly, the chosen dose, high when compared with studies estimating the dietary intake of the West Indian human population at up to 0.32µg/kg (Escal and Calbas studies 2003-2005) (Cornely, and Théodore, 2007; Merle, et al., 2008). The route of administration and duration of exposure also clearly differ from that of the human population. Our study also confirmed the missing capacity of mice to metabolize chlordecone in chlordecol, contrary to humans, requiring cautious interpretation of experimental data. However, mice were successfully used as a model of acute neurotoxicity found in humans, showing its relevance for studying toxic effects of chlordecone on neurotransmission.

The first experimental aim of this preliminary study was to identify possible early Parkinson’s-like neurological damage, since this has been previously reported with many other organochlorines in mice such as endosulfan (Wilson et al., 2014), heptachlor (Richardson et al., 2008) or dieldrin (Fleming et al., 1994) (Richardson et al., 2006). By comparison to the observed trend (∼15% decrease) of the number of dopaminergic neurons in the SNpc of mice in our study, a similar protocol with heptachlor, at a bi-weekly intraperitoneal dose of 7 mg/kg, revealed a selective dopaminergic loss (∼ 43%) in the SNpc and neuroinflammation in the ventral midbrain area, associated with subtle motor deficits revealed by functional tests (Hong et al., 2014). This is the only study showing dopaminergic loss by stereology among the rare studies in which adult mice were exposed to organochlorines by intraperitoneal route (Hatcher et al., 2007; Schuh et al., 2009; Hong et al., 2014), and to our knowledge, this has never been reported after oral exposures of adult mice so far. Interestingly, whereas a contamination of the human population through milk consumption was identified with heptachlor in Hawaii, an association with Lewy pathology, the cardinal feature of Parkinson’s disease, was recently reported (Ross et al., 2019). Among the dopaminergic markers examined by Western blot in our study, we noticed a VMAT2 decrease in the striatum of CLD-exposed mice. Whereas VMAT2 has been considered as protective against neurotoxic chemicals (Miller et al., 1999; Lohr and Miller, 2014; Lohr et al., 2016), a dose-dependent VMAT2 decrease has previously been reported in the striata of mice exposed by repeated intra-peritoneal injections of another organochlorine, methoxychlor (Schuh et al., 2009). These first data support the need of longer exposures of mice to question the possible occurrence of clearer dopaminergic deleterious effects.

Most importantly, our goal was then to provide the proof of principle for using functional ultrasound imaging in toxicology (toxico-fUS) in live animals, during a protocol of exposure to a pesticide. Using the fUSi method, we were indeed able to study the CLD effects at the end of the 2 months exposure. After visual stimulation of the animals, this approach enabled us to observe a significant decrease in cerebral blood volume (CBV) in the anterolateral and posteromedial areas of the visual cortex. It should be noted that all other cortical areas associated with vision showed a decrease in hemodynamic response, but this variation was non-significant. Patients with Parkinson’s disease frequently report problems with visual ability (Bowen et al., 1972; Fénelon et al., 2000; Archibald et al., 2011) as described by Weil et al: “Visuo-perceptual problems are well described in dementia with Lewy bodies and in Parkinson’s disease dementia, but there is growing evidence pointing to changes in visual processing earlier in the disease course” (Weil et al., 2016). Nevertheless, significantly reduced CBV changes only in cortical visual areas (and not other areas involved in visual processing such as in superior colliculus, where only a trend was found) suggest that the cortex itself is the most impaired region in this decreased response, with possible minor impairment of other regions. It remains to be understood to which extent these observations could remind visual deficits reported in CLD-exposed children (Desrochers-Couture et al., 2022) (Saint-Amour et al., 2020). The neuroanatomical basis of these deficits in both humans and mice still remain to be determined.

To the best of our knowledge, no experimental studies have so far examined the long-term neurotoxic effects of CLD in mammals, although such studies are clearly needed. Notably, an atypical parkinsonism has been specifically described in the French West Indies (Caparros-Lefebvre et al., 2002). This disease has been putatively associated with high consumptions of *Annona muricata* fruit, which worsens disease severity and cognitive deficits (Cleret de Langavant et al., 2022), and a low educational level was also intriguingly identified as a major risk factor (Lannuzel et al., 2018). A complex etiology, which may involve several neurotoxins (Caparros-Lefebvre et al., 2006), cannot be excluded. In this context, of high interest in the field, Parrales-Macias recently demonstrated that exposure to CLD caused a dose-dependent loss of dopaminergic in mouse midbrain cultures, and, associated with locomotor behavior deficits, in *C. elegans* worms (Parrales-Macias et al., 2023). Cholinergic and serotoninergic neuronal cells were also affected by CLD in *C. elegans*, although to a lesser extent than dopaminergic neurons. In humans, results from the TIMOUN mother-child cohort in Guadeloupe showed that CLD exposure impaired behavior, and both motor and cognitive abilities (Dallaire et al., 2012; Boucher et al., 2013; Desrochers-Couture et al., 2022; Oulhote et al., 2023).

In this context, our preliminary study in mice reinforce the hypothesis of a possible CLD involvement in parkinsonisms or/and neurodevelopmental disorders. This needs to be further investigated in mice thanks to longer, as well as oral or/and perinatal CLD exposures. Several studies also showed that exposures to organochlorines, at least during development, rendered dopaminergic neurons more vulnerable to subsequent neurotoxic challenges later in life ((Richardson et al., 2006) (dieldrin); (Boyd et al., 2023) (dieldrin) ; (Jia and Misra, 2007) (endosulfan) ; (Wilson et al., 2014) (endosulfan) ; (Richardson et al., 2008)). However, together with additional experimental research, our data emphasize the crucial need to follow parkinsonian syndromes in the French West Indies, which have still received little attention so far.

## Acknowledgements

fUS experiments were performed on CERMEP – Imagerie du vivant, Bron, F-69677, France, imaging facilities.

Jade Ruard (Lyon Neuroscience Research Center) is thanked for her technical help (cryosectioning).

We acknowledge the contribution of SFR Biosciences (Universite Claude Bernard Lyon 1, CNRS UAR3444, Inserm US8, ENS de Lyon) for the help of the staff of LYMIC-PLATIM especially Jacques Brocard and Elodie Chatre for assistance and advice with confocal scanner CQ1 used for stereology approach.

For the purpose of Open Access, a CC-BY 4.0 public copyright licence has been applied by the authors to the present document and will be applied to all subsequent versions up to the Author Accepted Manuscript arising from this submission.

## CRediT authorship contribution statement

Conceptualization: BV, GLB, TB, JV Formal analysis: LV, BV, FC, JV Funding acquisition: GLB, TB

Investigation: JV, FC, MV, BV, GLB, CD, MH, CA, EM

Methodology: BV, JV, MH Project administration: JV Resources: AD, DG, LL Software: LV, BV, EM Validation: FC, BV, GLB, CD

Visualization: FC, BV, GLB, CD, JV, MH Writing – original draft: JV

Writing – review & editing: FC, BV, TB, GLB, MH

## Declaration of Competing Interest

The authors declare that they have no known competing financial interests or personal relationships that could have appeared to influence the work reported in this paper.

## Supplementary information

### Isotopic dilution liquid chromatography–tandem mass spectrometry for quantification of CLD in brain tissue

#### Sample extraction procedure development

Different sample preparation protocols were tested based on the former dedicated methods implemented in the laboratory for food samples and bovine serum (Lavisson-Bompard et al. 2021b). For those methods, Quechers extraction is performed taking into account the amount of water in the sample : so after ACN addition and extraction, water is removed from the extract by demixion with the addition of the Qechers salts. Those methods are based on larger sample amount than in this study: typically 5 g for food matrices and 1 mL for bovine serum. Brains samples being much smaller, and grounded with very limited volume of distilled water, it was decided not to study both water addition and demixion but to test simple ACN extraction and dispersive SPE purification.

At first attempts, ACN extraction, with different solvent volumes were performed to determine the concentration levels expected in the brain sample, on a randomly selected contaminated sample (3) to adjust the sample/ACN ration regarding the sensitivity of the LC/MS² analytical method. A final extraction of the samples with 20 mL of ACN was selected so that the corresponding signal intensity for CLD being in upper middle range of the calibration curve levels to be validated.

The addition of dispersive SPE after ACN was also studied to reduce matrix effects. Comparison were performed in triplicate with and without dSPE, on spiked blank brain samples. After centrifugation and filtration of the supernatants, yields were close below 1 % with dSPE whether they were 96% + /-1%.

The very low recovery observed with dSPE can be explained by the size of the sample regarding usual more heavy ones (5g) : brains sample being of very low amount, the quantity of extraction sorbent is to high regarding the matrix content and thus CLD and CLDOH are also absorbed and retained on the dSPE. The use of dSPE was therefore abandoned.

So the final sample preparation consisted in a simple solvent extraction with 20 mL ACN, followed by the filtration of 2 mL of supernatant on syringe filter before HPLC analysis.

#### Method validation

Method validation and quality assurance/quality control. For method validation, the tolerances were set according SANTE guidance on retention time (+/-0.1 min relative deviation with regards to the standard), transitions rations (+/-30 % of the average calibrations standards in the extract) and recoveries (70-120% with repeatability bellow 20 %). Accuracy profiles were selected as statistical descriptors to determine and accept the gobal performances of the method, ie adequacy of the selected calibration function, accuracy and uncertainty of the method (Mermet and Granier, 2012) as per standard NF V03-110: 2010 (AFNOR, 2010). Probability β was set to 80% the risk of results lying outside the limits is below 20% on average. The expanded uncertainty (k=2) of the method was estimated taking into account both the standard deviation of the tolerance interval and the estimation of the bias.

The validation was performed on five series of blank mice samples, spiked at 4 concentration levels from 3 to 500 µg.kg-1 fresh weight (fw), each series was prepared on different days and each extraction / analyses performed in duplicate.

#### Range determination

The range levels were chosen according to the estimated concentration of CLD and CLD-OH in the brain on test samples portions. The chosen range varies from 0.5 ng/ml to 75 ng/ml. Since the collected brain samples are of different weight, it was decided to validate the method regarding the final diluted concentration of CLD and CLD-OH is solution (ng/mL) and to convert afterwards the results in mg/kg of brain. The accuracy profiles obtained for CLD and CLD-OH are presented in **supplementary figure S1**. In order to provide variability in the analyses, the method was performed in on-going mode, including all the suited brains regions in the series of validation.

The analytical procedure is validated for both CLD and CLD-OH form the experimental limit of quantification (LoQ) of 0.045 ng/mL, up to 75 ng/mL with respect to the SANTE guidelines requirements. CLD-OH accuracy profile is larger the CLD one: this is due to a larger dispersion of the recoveries with CLD-OH, since the internal standard, CLD-13C10, cannot exactly correct the matrix effect on CLD-OH.

The expanded uncertainty of the method was estimated taking into account the standard deviation of the tolerance interval: It was checked and set at 20%, for CLD and 25 % for CLD-OH corresponding to the maximum estimated expanded uncertainty on the studied concentration range.

---

**Supplementary figure S1.**
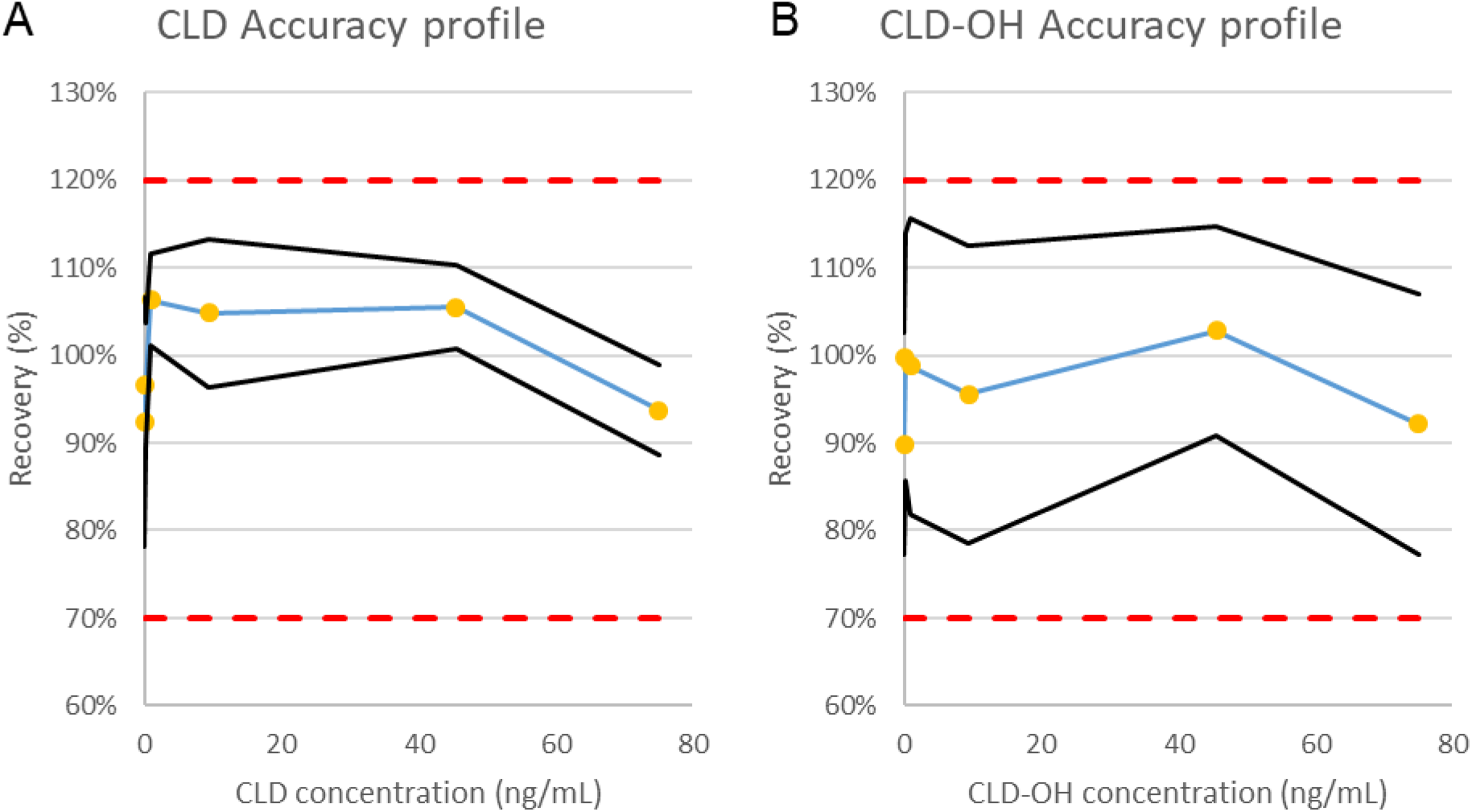
Accuracy profiles definition. The accuracy profiles were defined for CLD (**A)** and CLD-OH (**B**) form the experimental limit of quantification (LoQ) of 0.045 ng/mL, up to 75 ng/mL. Higher and lower acceptance limit were defined taking into account the width of the recovery and CLD-OH accuracy profile and the standard deviation of the tolerance interval.

**Supplementary figure S2.**
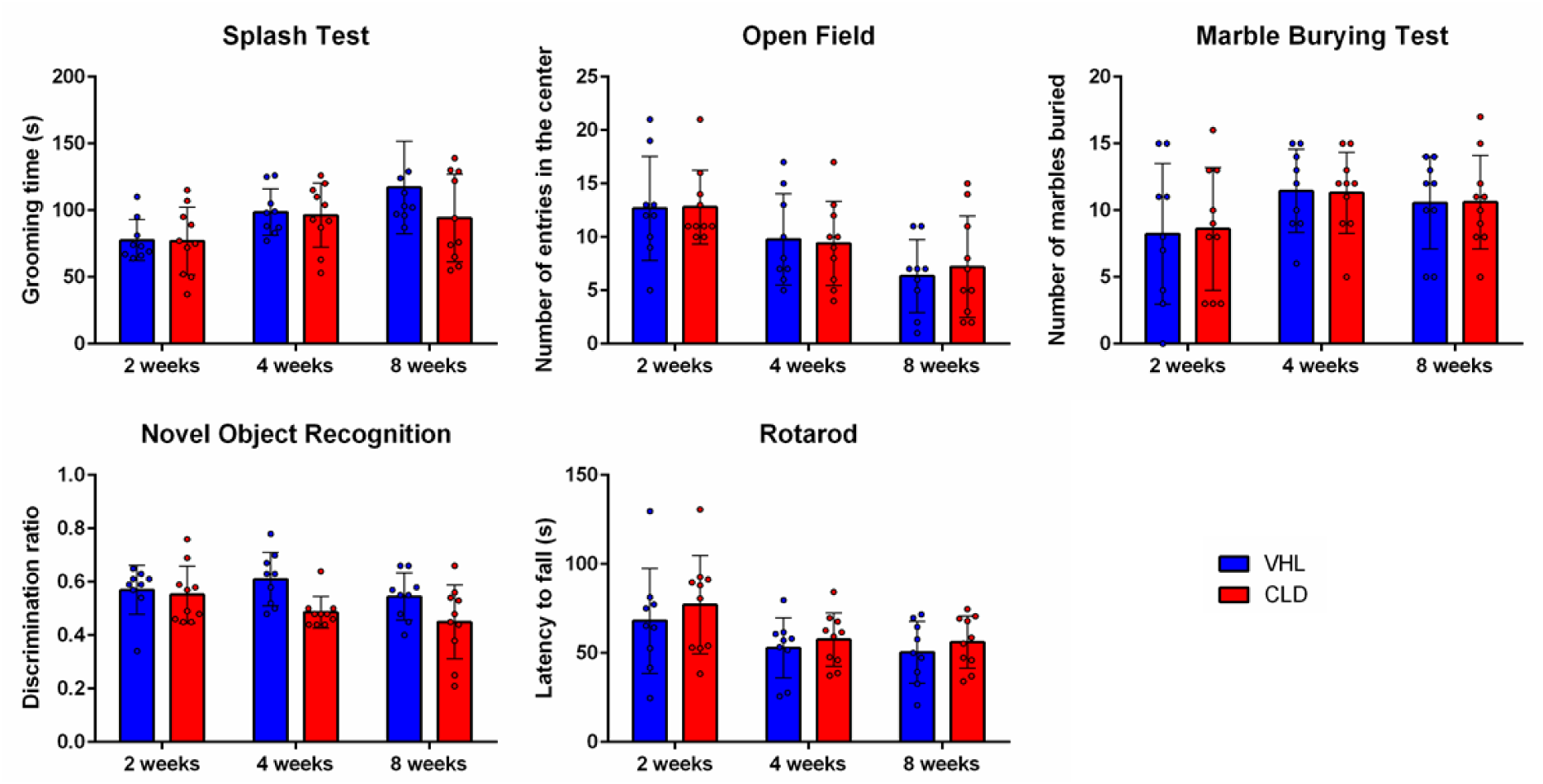
Behavioral assessments reveal no significant anxio-depressive or cognitive deficits following chlordecone exposure. Bar graphs depict behavioral performance in VHL (vehicle control, n = 9) and CLD (chlordecone-exposed, n = 10) mice at 2, 4, and 8 weeks of exposure. (A) Splash test: Total grooming time (seconds) as an indicator of motivational drive and depressive-like behavior. (B) Open Field test: Number of entries into the center zone, reflecting anxiety-like behavior. (C) Marble Burying test: Number of marbles buried (≥ two-thirds covered), a measure of repetitive/anxiety-like behavior. (D) Novel Object Recognition (NOR) test: Discrimination ratio calculated as time spent exploring the novel object divided by total exploration time, used to assess recognition memory. (E) Rotarod test: Average latency to fall (seconds) from the accelerating rotating rod over three trials, used to assess motor coordination. Data are expressed as mean ± SD.

**Supplementary figure S3.**
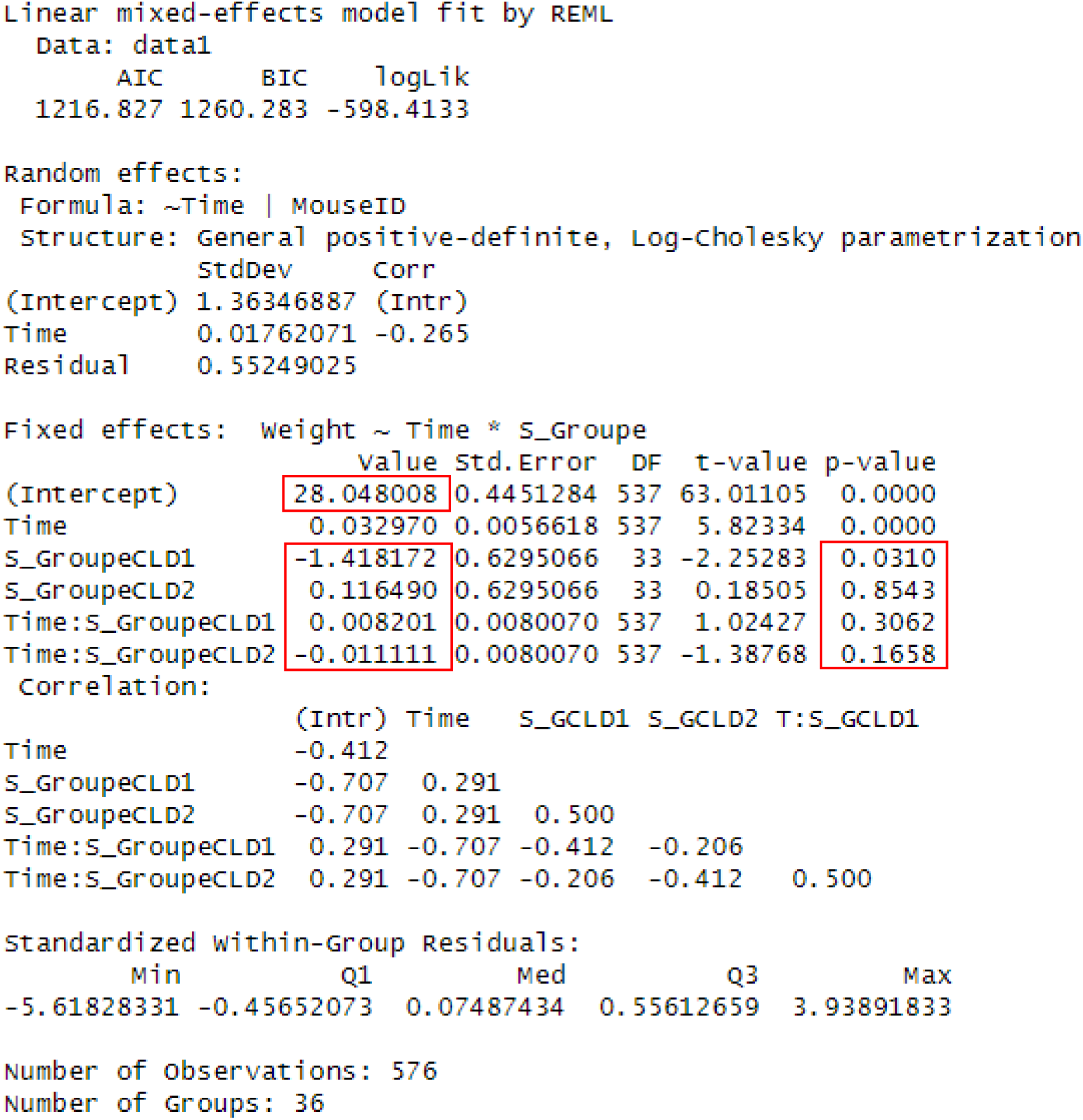
Evolution of the mice mass measured over time modeled using mixed effects models. In this set evolution of bodyweight of CLD1, CLD2 and VHL1 were compared. This figure represents results from model 1 with VHL1 as reference. We compared the average mass of each group at the start of the experiment (Intercept) and the progression of mass over time (Time). Red squares represented value commented in the article.

**Supplementary figure S4.**
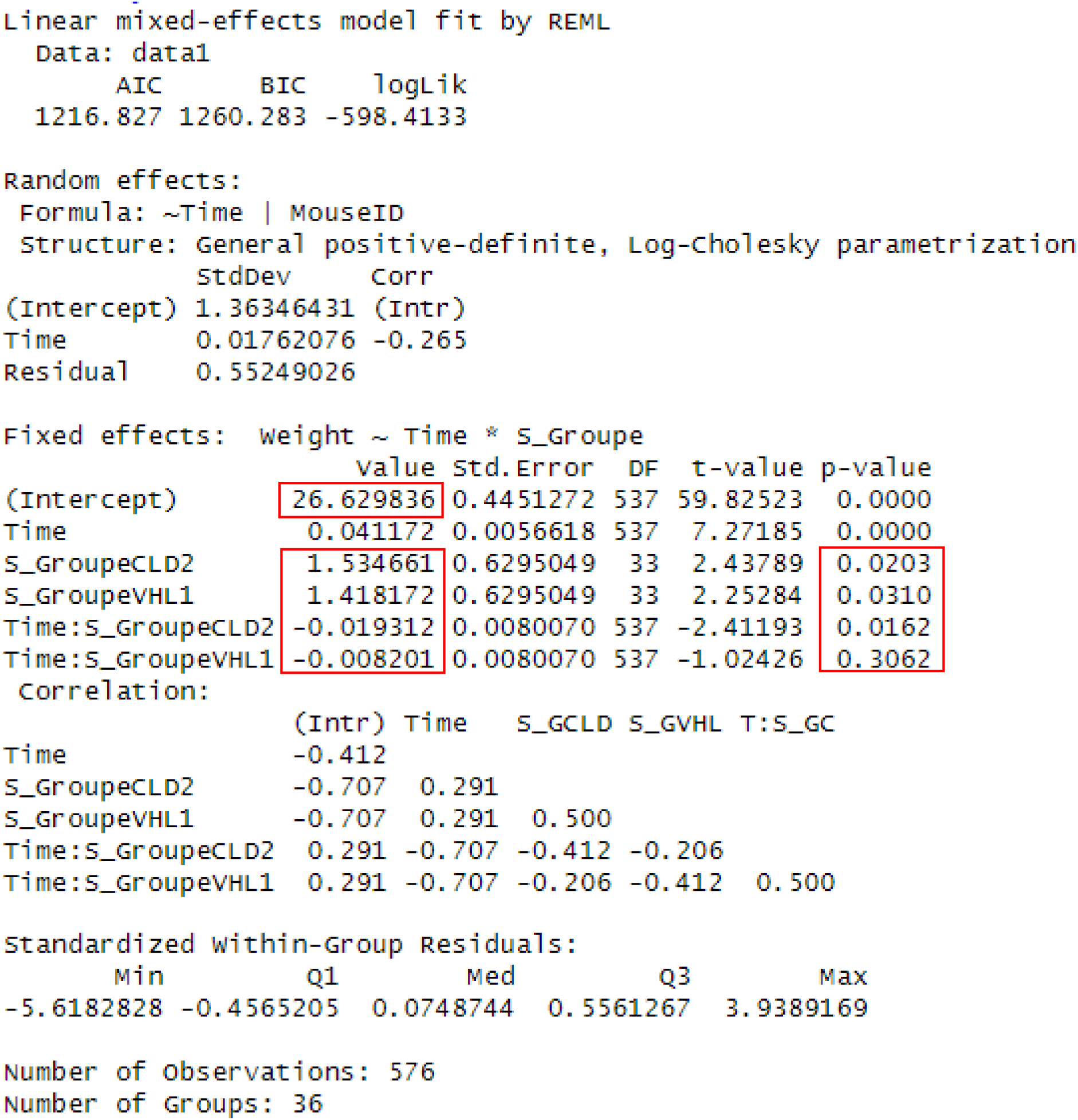
Evolution of the mice mass measured over time modeled using mixed effects models. In this set evolution of bodyweight of CLD1, CLD2 and VHL1 were compared. This figure represents results from model 1 with CLD1 as reference. We compared the average mass of each group at the start of the experiment (Intercept) and the progression of mass over time (Time). Red squares represented value commented in the article.

**Supplementary figure S5.**
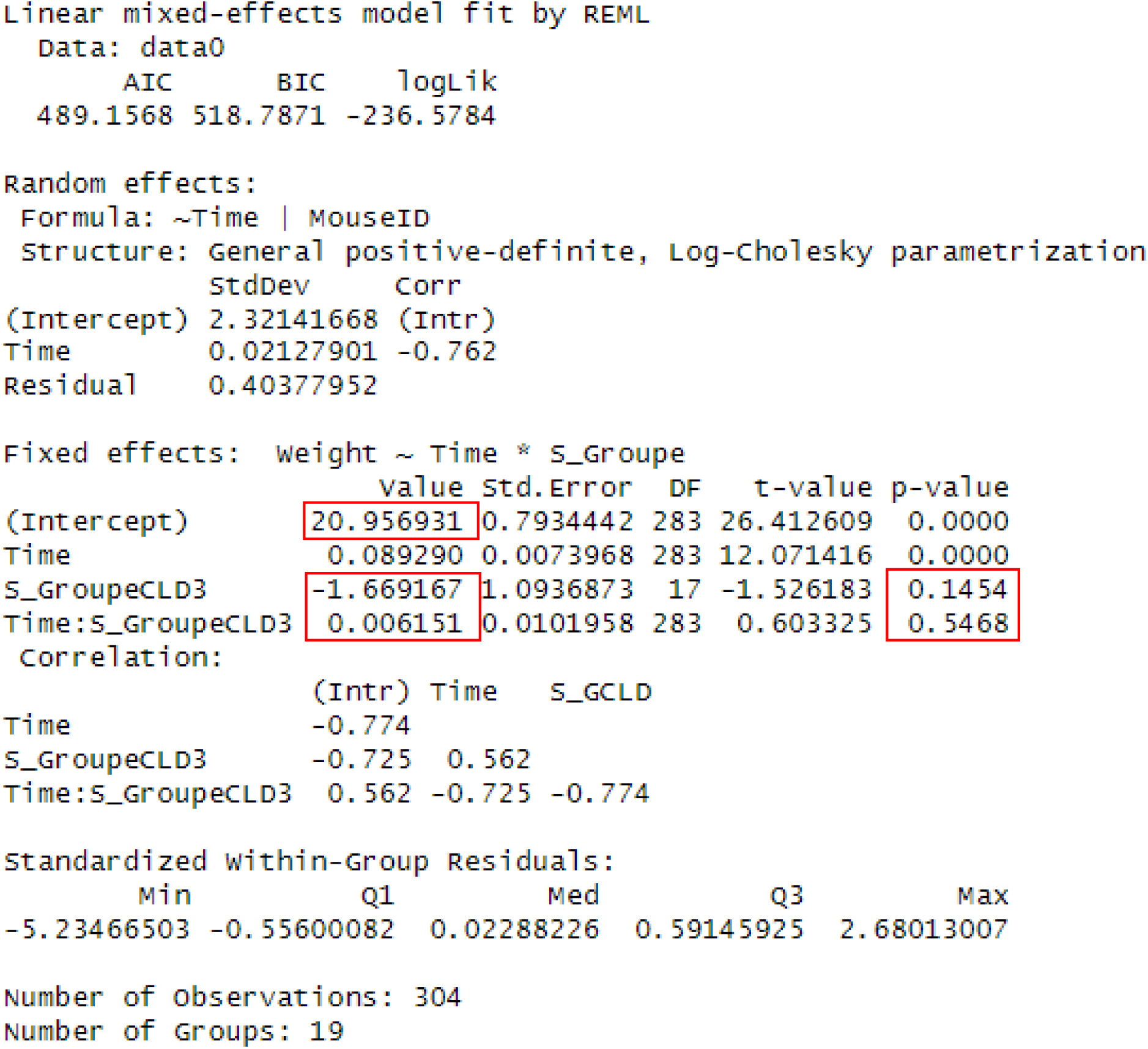
Evolution of the mice mass measured over time modeled using mixed effects models. In this set evolution of bodyweight of CLD3 and VHL2 were compared. This figure represents results from model 2 with VHL2 as reference. We compared the average mass of each group at the start of the experiment (Intercept) and the progression of mass over time (Time). Red squares represented value commented in the article.

**Supplementary figure S6.**
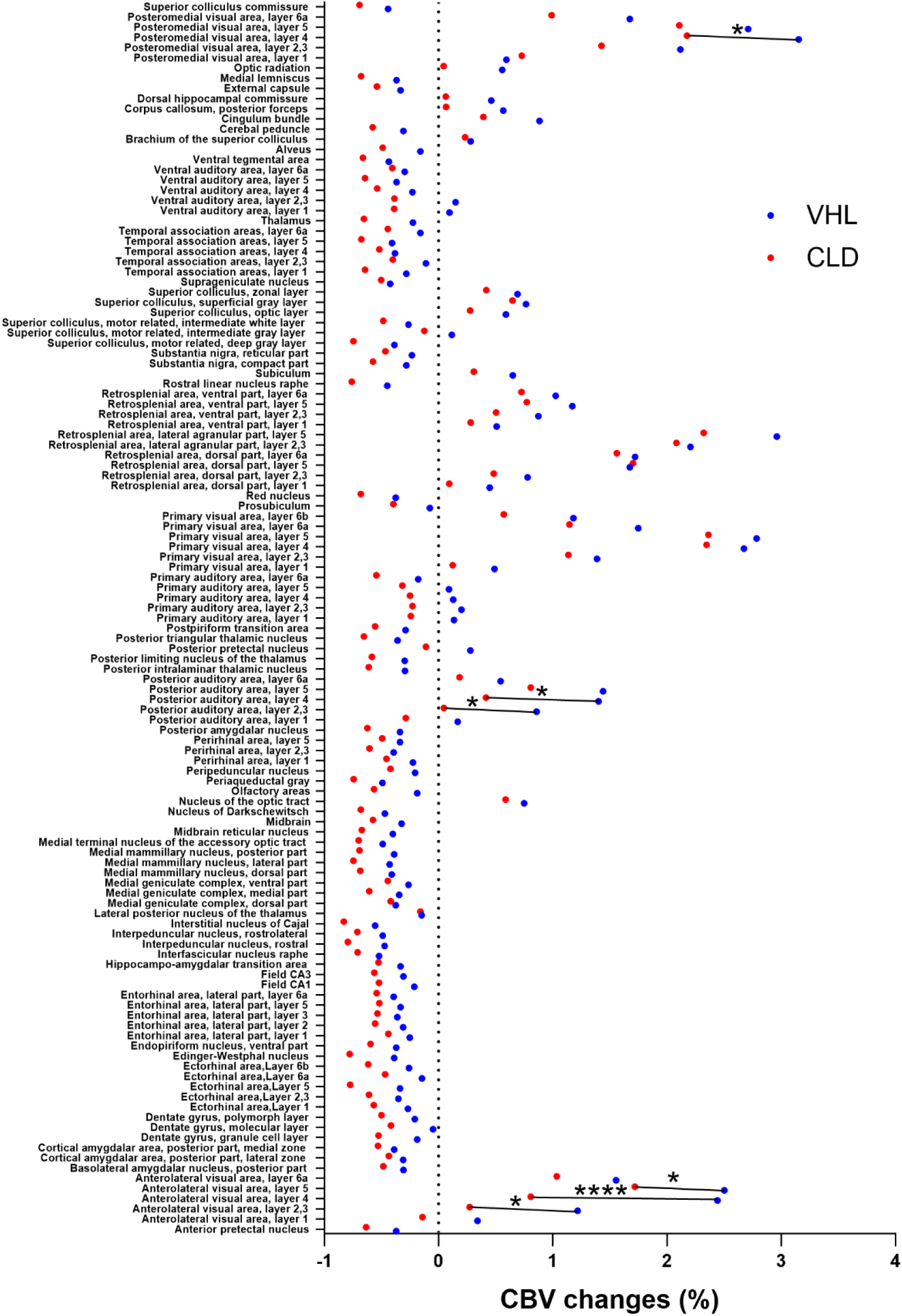
Alteration of response to visual stimulation following chlordecone exposure detected by fUSi. Points depict the mean changes of CBV following visual stimulation compared to baseline for each group (VHL in blue, n=9; CLD in red, n=10), in the regions of the Allen atlas at the corresponding coronal slice. Significant differences between VHL and CLD in specific regions are shown by lines connecting the two groups (two-way ANOVA followed by FDR method; *q<0.1, ****q<0.0001).

## Notes

### Competing Interest Statement

The authors have declared no competing interest.

### Summary of Updates

One author added Statistical analysis of mice body weight have been added Figure 1 revised

